# DNA passes through cohesin’s hinge as well as its Smc3-kleisin interface

**DOI:** 10.1101/2022.05.30.494034

**Authors:** James E Collier, Kim A Nasmyth

## Abstract

The ring model (Haering et al. 2002) proposes that sister chromatid cohesion is mediated by co-entrapment of sister DNAs inside a tripartite cohesin ring created by a pair of rod-shaped proteins (Smc1 and Smc3) whose two ends are connected through dimerization of their hinges at one end and by association of their ATPase domains at the other end with the N- and C-terminal domains of a kleisin subunit (Scc1). The model explains how Scc1 cleavage triggers anaphase (Uhlmann, Lottspeich, and Nasmyth 1999) but has hitherto only been rigorously tested using small circular mini-chromosomes in yeast, where crosslinking the ring’s three interfaces, creating a covalent circular molecule, induces catenation of individual sister DNAs (Haering et al. 2008; Srinivasan et al. 2018). If the model applies to real chromatids, then the ring must have a DNA entry gate essential for mitosis. Whether this is situated at the Smc3/Scc1 (Murayama and Uhlmann 2015; Murayama et al. 2018) or Smc1/Smc3 hinge (Gruber et al. 2006) interface is an open question. Using an in vitro system (Collier et al. 2020), we show that cohesin in fact possesses two DNA gates, one at the Smc3/Scc1 interface and a second at the Smc1/3 hinge. Unlike the Smc3/Scc1 interface, passage of DNAs through SMC hinges depends on both Scc2 and Scc3, a pair of regulatory subunits necessary for entrapment in vivo (Srinivasan et al. 2018). This property together with the lethality caused by locking this interface but not that between Smc3 and Scc1 in vivo (Gruber et al. 2006) suggests that passage of DNAs through the hinge is essential for building sister chromatid cohesion. Passage of DNAs through the Smc3/Scc1 interface is necessary for cohesin’s separase-independent release from chromosomes (Chan et al. 2012) and may therefore largely serve as an exit gate.

## Introduction

The sister chromatid cohesion essential for mitosis and meiosis is mediated by a pair of rod-shaped SMC proteins (Smc1 and Smc3) joined together through an interaction between hinge domains at one end. The interconnection by a kleisin subunit (Scc1) of their ATPase domains at the other end creates a ring-like structure within which it is proposed sister DNAs are entrapped during DNA replication (Haering et al. 2002). Two approaches have hitherto been used to test the ring model. The first has been a method to induce thiol-specific chemical crosslinks within the three interfaces of SMC-Kleisin (S-K) rings. An early version of this approach showed that covalent circularization by BMOE of a version of cohesin containing cysteine pairs within the ring’s interfaces is sufficient to cause catenation of small circular sister DNAs that are otherwise not inter-twined, hence proving their co-entrapment (Haering et al. 2008). Furthermore, analysis of a wide variety of strains carrying different cohesin mutations has subsequently confirmed that catenation in this manner of sister minichromosome DNAs correlates with the ability of yeast cells to proliferate (Srinivasan et al. 2018).

The second approach has been to elucidate the mechanism by which DNAs enter S-K rings. The logic being that only when we have understood this mechanism and found it to operate inside cells could we be certain that entrapment does indeed form the basis of cohesion. The initial goal was to establish which of the cohesin rings’ three interfaces must open up to let in DNA, in other words to identify cohesin’s DNA entry gate. The finding that inter-connection of Smc1 and Smc3 hinges using rapamycin (when FKBP12 and FRB were inserted into small loops within the Smc1/3 hinges) blocked cohesion establishment while fusion of Smc3 or Smc1 with Scc1 did not do so led to the proposal that if cohesin has a unique essential DNA entry gate, then it must be at the hinge interface (Gruber et al. 2006). However, these experiments merely showed that a modification predicted to hinder hinge opening blocks establishment of cohesion, which is not the same as proving that DNAs actually enter via this interface. Besides which, the conclusion that the hinge is a DNA entry gate is not universally accepted (Murayama and Uhlmann 2015; Murayama et al. 2018). Thus, despite being crucial for understanding how cohesion is established, the location of cohesin’s DNA entry gate remains unresolved. To break this impasse, we recently developed an in vitro assay to measure entrapment of DNAs within S-K rings, using the same technique used in vivo, namely catenation of circular DNAs by cysteine-substituted cohesin rings chemically circularized using BMOE (Collier et al. 2020). S-K DNA entrapment in vitro is stimulated by Scc2 and depends on ATP and Scc3 but not on Pds5 or Wapl, reflecting the properties of S-K entrapment in vivo (Srinivasan et al. 2018), suggesting the reaction is physiologically relevant. Using this system, we now describe experiments that show definitively that cohesin possesses two gates through which DNAs pass, at least in vitro, one at the Smc3/Scc1 interface and a second at the Smc1/3 hinge. We also describe a series of topological assays suggesting that the first step is passage of DNAs between cohesin’s ATPase heads and their enclosure by Scc2 in the lower half of an SMC compartment bounded by hinges and heads engaged by ATP.

### Covalent closure of cohesin ring interfaces

Cohesin’s DNA gate(s) could in principle be identified merely by observing the process of entrapment in real time. Because this is not at present technically possible, we instead covalently sealed each ring interface shut in a manner orthogonal to the thiol-specific crosslinking protocol that we use to measure entrapment (Collier et al. 2020). This approach allows us to seal potential DNA gates prior to the entrapment process. If DNA entered through a single gate, then sealing should block entrapment. Though such a result would be consistent with the gate being used for entrapment, it would not directly demonstrate passage, as the sealing process could in principle also interfere with passage through a different gate, by somehow altering the latter’s conformation. Crucially, a failure to block entrapment by sealing a single ring interface would imply that the interface in question is either not a gate or that that it is not the sole entry gate. Through a striking result, blocking entry would therefore permit only weak conclusions. A more rigorous approach would be to seal two out of the ring’s three interfaces, leaving only a single potential gate. In this case, continued entrapment would demonstrate that DNA must have passed through the sole remaining interface.

To seal the Smc3/Scc1 interface, we expressed a fusion protein in which the N-terminus of Scc1 is connected by a short linker to the C-terminus of Smc3 (S3 fusion; Figure 1A, Figure 1 - supplement 1A). A similar approach was used connect Scc1’s C-terminus to Smc1’s N-terminus, thereby sealing the Smc1/Scc1 interface (S1 fusion). Smc1 and Smc3 hinges cannot be connected using this approach because they are located within the middle of their polypeptides. In this case, we achieved highly efficient covalent closure using an isopeptide bond created between a spytag and a spycatcher domain (Li et al. 2014) inserted into loops on the surface of Smc1 and Smc3 hinges respectively (Hinge fusion; Figure 1A, Figure 1 - supplement 1A & B). We next created three types of cohesin complexes composed of a single polypeptide in which two out of three interfaces were covalently sealed, namely a Smc3-Scc1-Smc1 (S3 S1 fusion) fusion containing a wild type hinge interface, a Smc3-Scc1 fusion whose hinge was connected to that of Smc1 using a spytag-spycatcher pair (S3 Hinge fusion), and a Scc1-Smc1 fusion whose hinge was similarly connected to that of Smc3 (S1 Hinge fusion). Lastly, we created a covalently circular ring in which all three interfaces were sealed (Circular cohesin). Remarkably, all seven types of cohesin rings produced abundant and stable complexes (Figure 1B & C). Some had modestly reduced rates of ATP hydrolysis, with circular cohesin having the greatest defect (~ 60 % that of WT cohesin; Figure 1 - supplement 1C).

**Figure 1.**
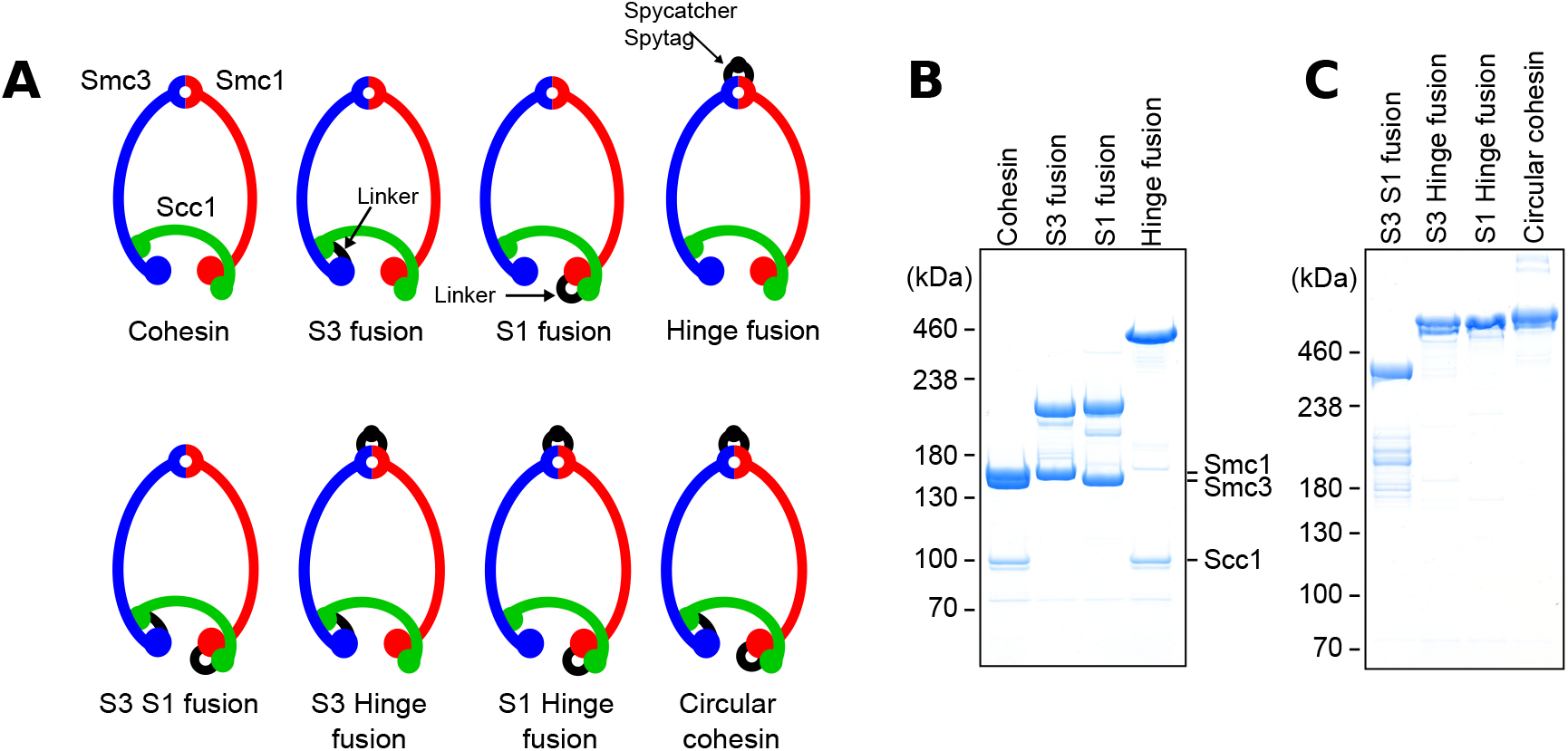
Covalent closure of cohesin’s interfaces. (A) SMC-kleisin rings, showing their three interfaces. Those connected by a covalent linkage are marked in black. (B) Coomassie stain of purified cohesin with either the Smc3/Scc1 (S3 fusion), Scc1/Smc1 (S1 fusion), or hinge interfaces (Hinge fusion) covalently sealed. (C) Coomassie stain of purified cohesin with two or three interfaces covalently sealed.

**Figure supplement 1.**
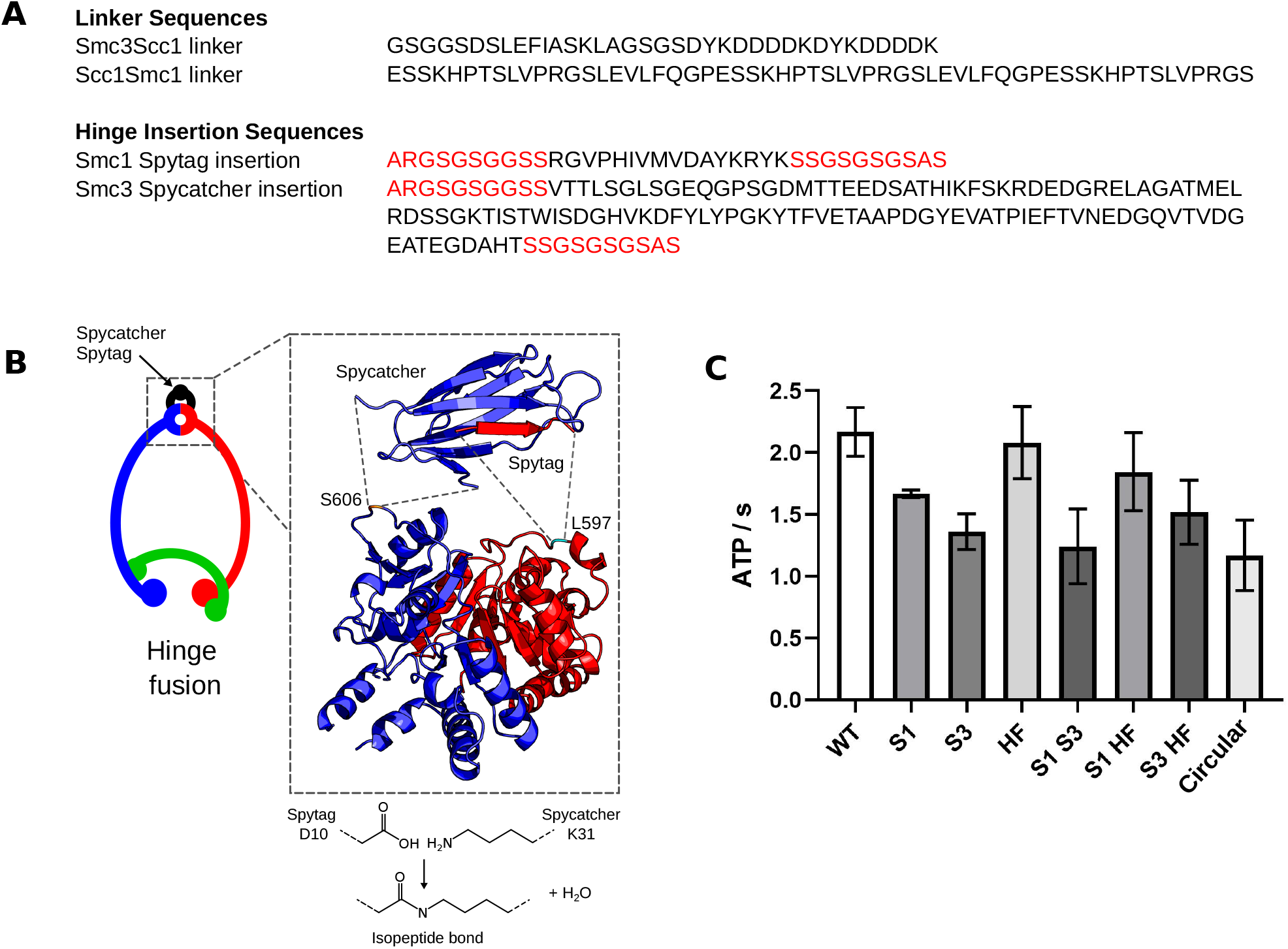
Covalent closure of cohesin’s interfaces. (A) Top; The polypeptide sequences used to fuse Scc1 to Smc1 and Smc3. Bottom; polypeptide sequences of the spytag and spycatcher inserted into Smc1 (after residue L597) and Smc3 (after residue S606), respectively. Linker sequences in red and spytag/spycatcher sequences in black. (B) Composite model (4MLI & 2WD5) of the spytag and spycatcher inserted into the Smc1 and Smc3 hinge domains, respectively. An isopeptide bond forms between D10 of the spytag and K31 of the Spycatcher. (C) ATPase data for cohesin constructs with interfaces covalently sealed compared to WT carried out in the presence of Scc2, Scc3, and DNA.

### DNA passes through the Smc1/Smc3 hinge as well as the Smc3/Scc1 interface

We next tested the ability of these different fusion proteins to entrap circular plasmid DNA in vitro (Figure 2 - supplement 2A; Collier et al. 2020). As expected from in vivo results (Gruber et al. 2006; Srinivasan et al. 2018), DNAs were entrapped by the cohesin rings containing either the Smc3-Scc1 or Scc1-Smc1 fusions. More surprising, they were also entrapped by rings with the Hinge fusion (Figure 2A). It is difficult to make direct comparisons between the efficiencies as entrapment by each version depends on different cysteine pair combinations, whose crosslinking by BMOE can vary. We can nevertheless conclude that DNA enters the cohesin ring via at least two different gates. To identify these, we measured entrapment by rings with two interfaces sealed, whereby DNA can only enter through the one remaining open interface. These assays revealed ATP-dependent DNA entrapment by cohesin rings containing the Smc3-Scc1-Smc1 fusion (S3 S1 fusion) as well as rings containing the Scc1-Smc1-Hinge fusions (S1 Hinge fusion; Figure 2B). However, DNA was not entrapped by a complex with the Smc3-Scc1-Hinge fusions (S3 Hinge fusion). These observations demonstrate that DNA entrapment arises by passage through the hinge as well as through the Smc3/Scc1 interfaces. Importantly, no passage occurs through the Smc1/Scc1 interface, at least when the other two gates are covalently sealed.

### Scc2 is required for DNA passage through the hinge but not through the Sm3/Scc1 interface

Entrapment within S-K rings in vivo normally depends on both Scc2 and Scc3. We therefore addressed whether hinge or Smc3/Scc1 gate passage in vitro shares this property. Entrapment by the Smc3-Scc1-Smc1 fusion was abolished by omission of either regulatory protein (Figure 2C). Entrapment via the hinge in vitro therefore resembles in vivo entrapment. This suggests that the reason why cohesion establishment is abolished by linkage of Smc1 and Smc3 hinges containing FRB and FKBP12 respectively using rapamycin (Gruber et al. 2006) is because passage of DNA through the hinge is an essential step. Interestingly, the strict dependence of hinge-mediated entrapment on Scc2 differs from entrapment by wild type S-K rings in vitro, which still occurs in the absence of Scc2, albeit at a reduced level (Figure 2 – supplement 2B; Collier et al. 2020). A simple explanation for this difference is that cohesin entraps DNA via both hinge and Smc3/Scc1 pathways in vitro and the Scc2-independent entrapment is due entirely to passage through the Smc3/Scc1 gate. If so, sealing the Smc3/Scc1 gate should abolish Scc2-independent S-K entrapment. As predicted, entrapment by rings containing the Smc3-Scc1 fusion depends on Scc2 as well as Scc3 (Figure 2D). In other words, Scc2-independent entrapment requires a Smc3/Scc1 gate that can be opened.

**Figure 2.**
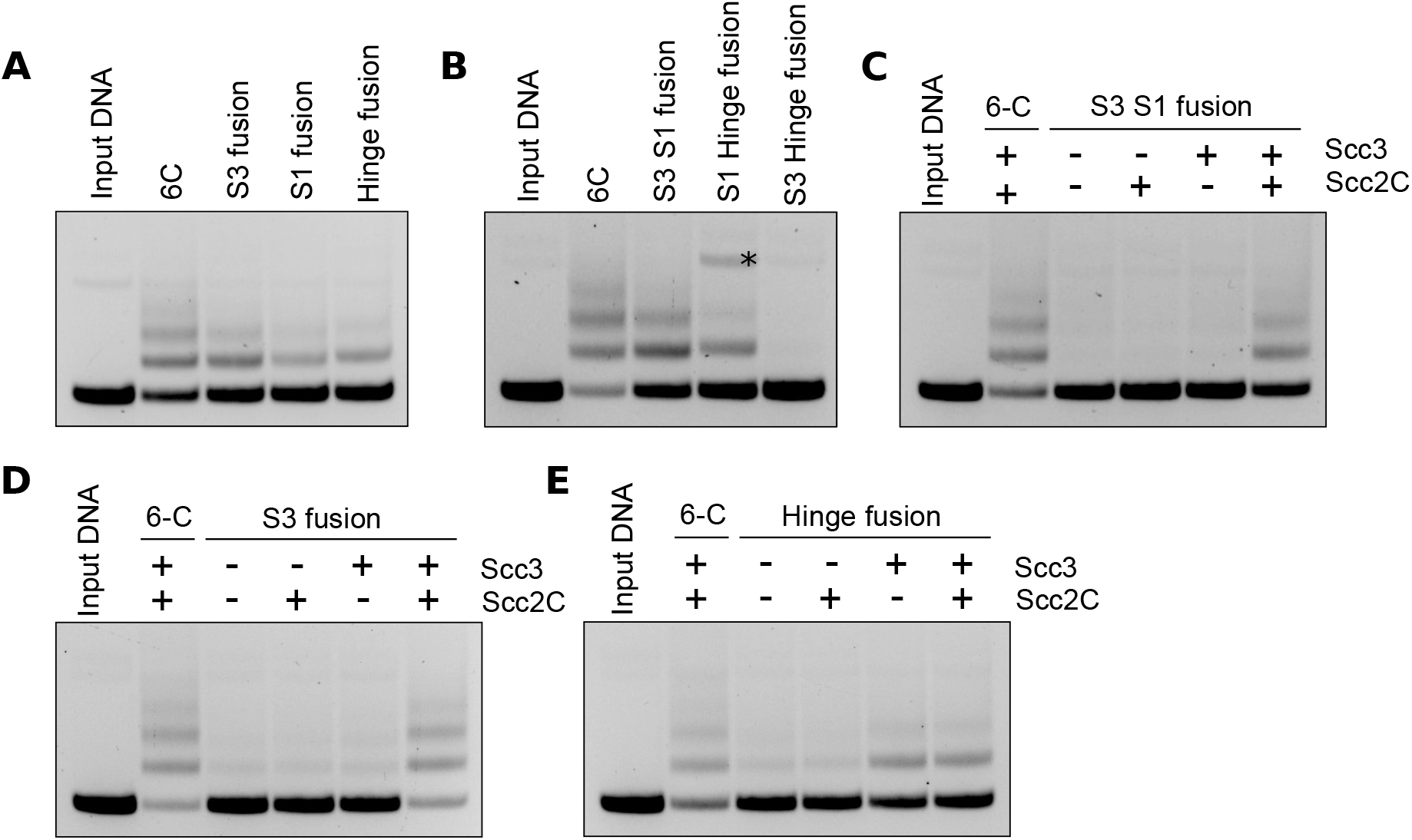
DNA passes through cohesin’s hinge and Smc3/Scc1 interfaces. (A) DNA entrapment assay comparing WT cohesin (6C) to constructs with either the Smc3/Scc1 (S3 fusion), Scc1/Smc1 (S1 fusion), or hinge (Hinge fusion) interfaces sealed in the presence of Scc2 and Scc3 after a 40 min incubation. (B) Entrapment of DNA by WT cohesin (6C) compared to constructs with either the Smc3-Scc1-Smc1 (S3 S1 fusion), Scc1-Smc1 and hinge (S1 Hinge fusion), or Smc3-Scc1 and hinge (S3 Hinge fusion) interfaces shut in the presence of Scc2 and Scc3 after a 40 min incubation. * = nicked DNA. (C) DNA entrapment comparing WT cohesin in the presence of Scc2 and Scc3 with the Smc3-Scc1-Smc1 fusion construct (S3 S1 fusion) in the presence or absence of Scc2 and Scc3 after a 40 min incubation. (D) DNA entrapment comparing WT cohesin (6C) in the presence of Scc2 and Scc3 with the Smc3-Scc1 fusion construct (S3 fusion) in the presence or absence of Scc2 and Scc3 after a 40 min incubation. (E) DNA entrapment comparing WT cohesin (6C) in the presence of Scc2 and Scc3 with the hinge fusion construct in the presence or absence of Scc2 and Scc3 after a 40 min incubation.

**Figure supplement 2.**
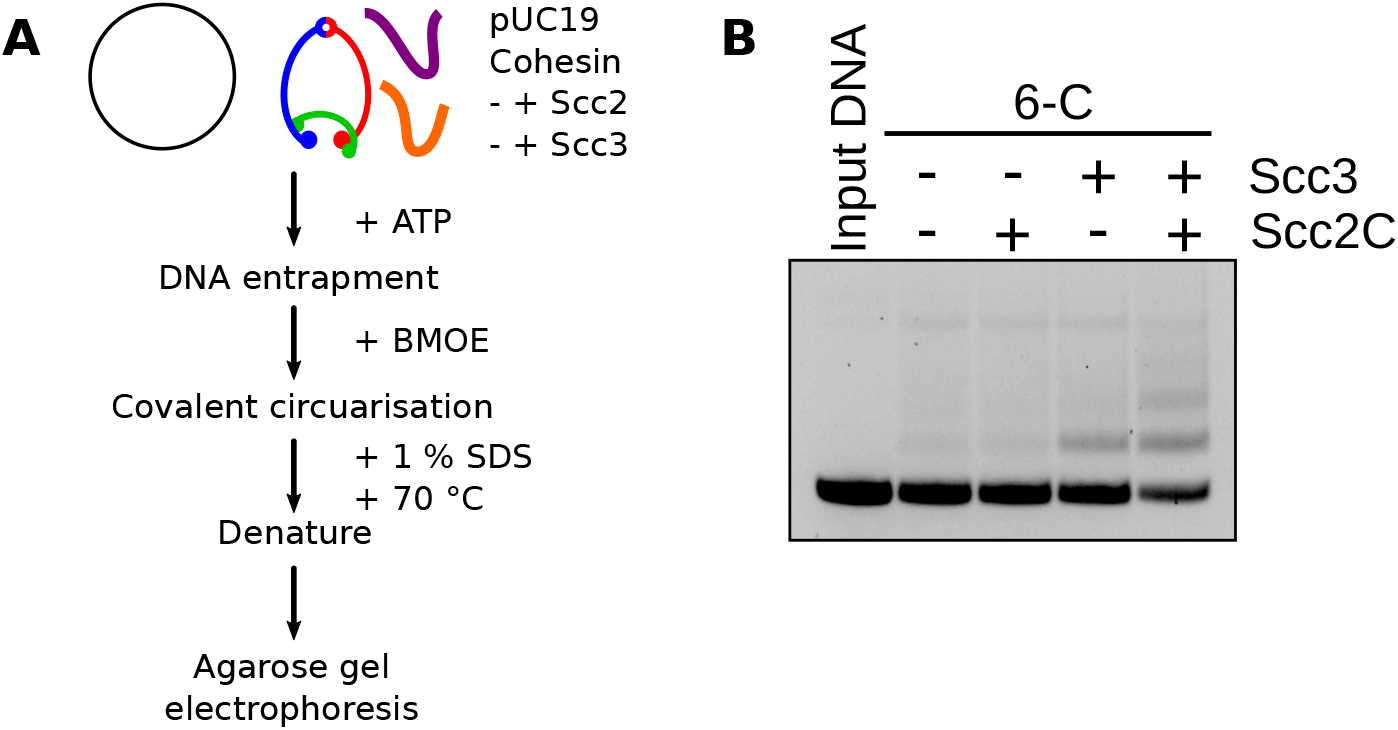
DNA passes through cohesin’s hinge and Smc3/Scc1 interfaces. (A) Schematic of the in vitro entrapment assay. (B) DNA entrapment for WT cohesin (6C) in the presence or absence of Scc2 and Scc3.

To address whether entrapment via the Smc3/Scc1 gate is affected by Scc2, we measured the effect of Scc2 and Scc3 on entrapment of DNA by cohesin whose hinge alone is fused and contains cysteine pairs within both SMC-Scc1 interfaces. Because DNA cannot pass through the Smc1/Scc1 interface, all entrapment by such cohesin must be via its Smc3/Scc1 interface. Entrapment by this construct depends on Scc3 but not Scc2 (Figure 2E). The dependence of DNA passage through the Smc3/Scc1 interface on Scc3 but not Scc2 in vitro resembles the activity that dissociates cohesin from chromosomes in vivo, a process dependent on Wapl and also blocked by the Smc3-Scc1 fusion (Chan et al. 2012). It is therefore possible that our in vitro experiments capture this process, albeit acting in reverse, as previously suggested (Murayama and Uhlmann 2015). In vivo, release not only does not require Scc2 but is actively blocked by it, at least in G1 cells (Srinivasan et al. 2019). Given that passage of DNA through the Smc3/Scc1 interface is not required for cell proliferation, for S-K entrapment in vivo, or even for cohesin’s stable association with the bulk of the genome, it is uncertain whether passage of DNA through this gate has any role in building cohesion in addition to its well documented role in mediating release.

### DNAs entrapped in SMC compartments in the absence of Scc3 are located between Scc2 and engaged heads

Given that passage of DNA through the hinge may be essential for building cohesion, the strict dependence of this process on Scc2 raises a question as to Scc2’s role. Reactions performed in the absence of Scc3 provide an important clue. Under these circumstances, Scc2 promotes rapid entrapment of DNA within cohesin’s SMC compartment (Figure 3 – supplement 3A), namely between the hinge and Smc1 and Smc3 head domains engaged in the presence of ATP (Collier et al. 2020). Crucially, this process is not accompanied by entrapment within S-K rings (Collier et al. 2020). Cryo-EM structures of DNA oligonucleotides bound to Scc2 and cohesin suggest that DNA trapped within SMC compartments by Scc2 binds simultaneously to Scc2 and a groove created by the engagement of Smc1 and Smc3 heads in the presence of ATP (PDB 6ZZ6). DNA associates with similar grooves above the engaged heads of condensin (Lee, Rhodes, and Lowe 2022), MukBEF (Burmann et al. 2021), and Rad50 (Kashammer et al. 2019), implying that this type of association is a highly conserved feature of SMC-like ATPase domains. In the case of cohesin, the DNA is actually “clamped” in a small compartment created by association of Scc2’s N-terminal and central domains bound to Smc3’s neck and head domains respectively (Collier et al. 2020; Higashi et al. 2020; Shi et al. 2020). Though the conditions under which these cryo-EM structures were obtained resemble those necessary for entrapment of DNA within the SMC compartment, namely both require Scc2, ATP, and DNA, but not Scc3 or ATP hydrolysis (Collier et al. 2020), we cannot be certain whether the two activities are truly synonymous. What is required is a crosslinking assay for DNA clamping comparable to the one used to measure SMC compartment entrapment.

We therefore designed a set of cysteine pairs within Scc2-SMC interfaces that could be crosslinked by BMOE if DNA were clamped in the manner observed in the cryo-EM structure (Figure 3 - supplement 3C). To this end, cysteines were introduced into the interfaces between Scc2 and the Smc1 head (Scc2T1281C Smc1E1102C), between Scc2 and the Smc3 head (Scc2E819C Smc3S72C), and between Scc2 and Smc3’s neck (Scc2D369C Smc3K1004C). As predicted by the cryo-EM structure, all three pairs enabled BMOE to crosslink Scc2 to SMC heads in the presence of ATP and DNA (Figure 3A - C). Crosslinking between Scc2 and either Smc1 or Smc3 head occurred in the absence of both ATP and DNA but was stimulated by ATP, an effect that was more pronounced for crosslinking between Scc2 and the Smc3 head. DNA also modestly increased crosslinking between both cysteine pairs, but only in the absence of ATP. In contrast, crosslinking between Scc2 and the Smc3 neck was strongly ATP dependent and enhanced by DNA (Figure 3C). These results suggest that Scc2 initially binds to the Smc1 head, subsequently binds the Smc3 head, and only binds the Smc3 neck efficiently upon the engagement of Smc1 and Smc3 heads in the presence of ATP and DNA.

**Figure 3.**
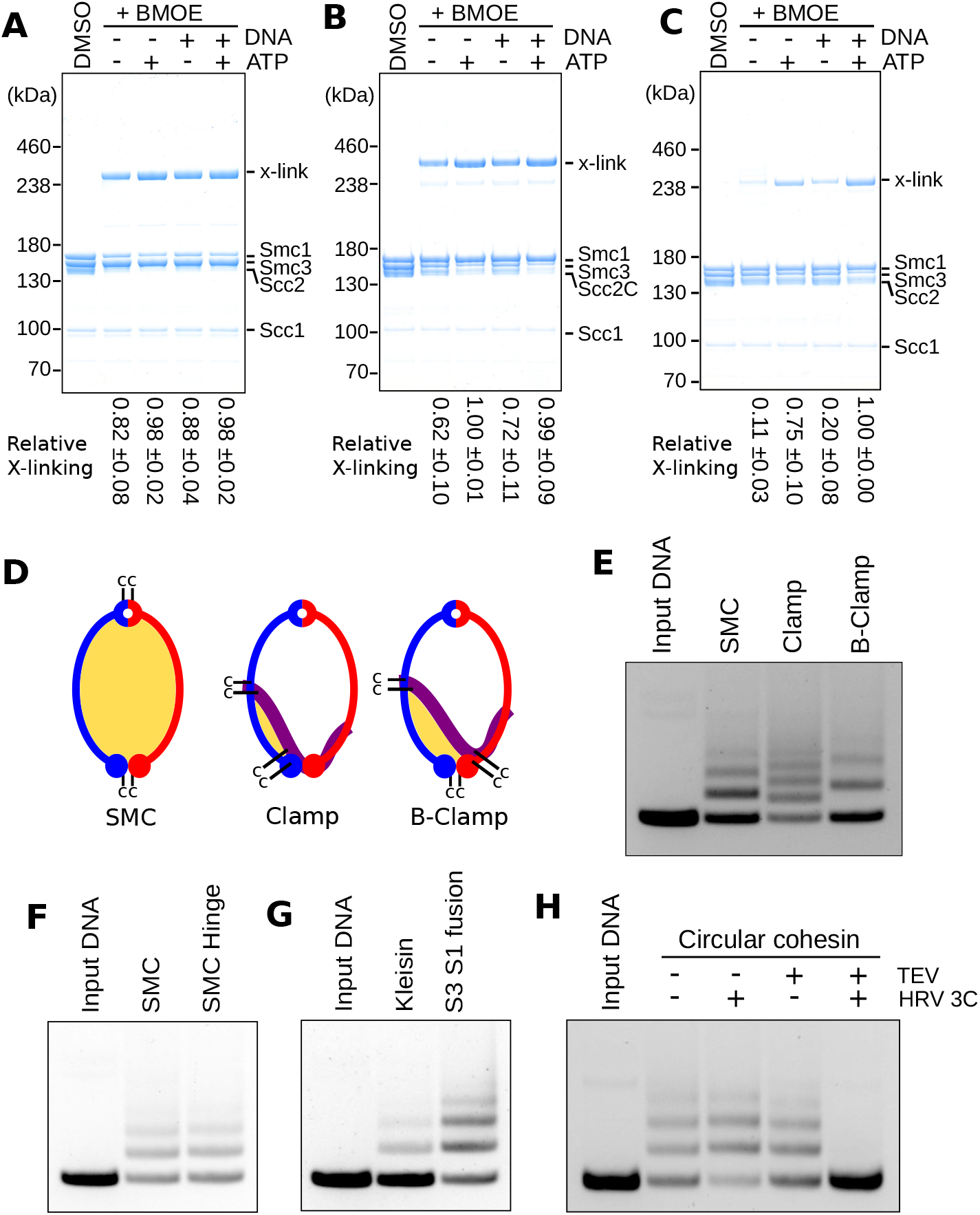
DNA passes through cohesin’s ATPase domains. (A) Crosslinking of Scc2 to the Smc1 head in the presence or absence of ATP and DNA. (B) Crosslinking Scc2 to the Smc3 head in the presence or absence of ATP and DNA. (C) Crosslinking Scc2 to the Smc3 neck in the presence or absence of ATP and DNA. (D) Models of cohesin showing either the SMC, Clamp, or below the clamp (B-Clamp) compartments, highlighted in yellow. For Clamp and B-Clamp compartments Scc2 is in purple. (E) Entrapment of DNA in either the SMC, Clamp, or below the clamp (B-Clamp) compartments, in the presence of Scc2 after a 2 min incubation. (F) Entrapment of DNA in the SMC compartment by cohesin with either a WT hinge (SMC) or with the hinge covalently fused (SMC Hinge) in the presence of Scc2 after a 2 min incubation. (G) Entrapment of DNA in the kleisin compartment by cohesin with either both kleisin interfaces open (Kleisin) or covalently closed (S3 S1 fusion) in the presence of Scc2 after a 2 min incubation. (H) Entrapment of DNA by covalently circular cohesin in the presence of Scc2 after a 2 min incubation. After crosslinking, BMOE was quenched by addition of DTT and then the samples were treated with TEV and/ or HRV 3C proteases and incubated at 24 °C for 30 min.

**Figure supplement 3.**
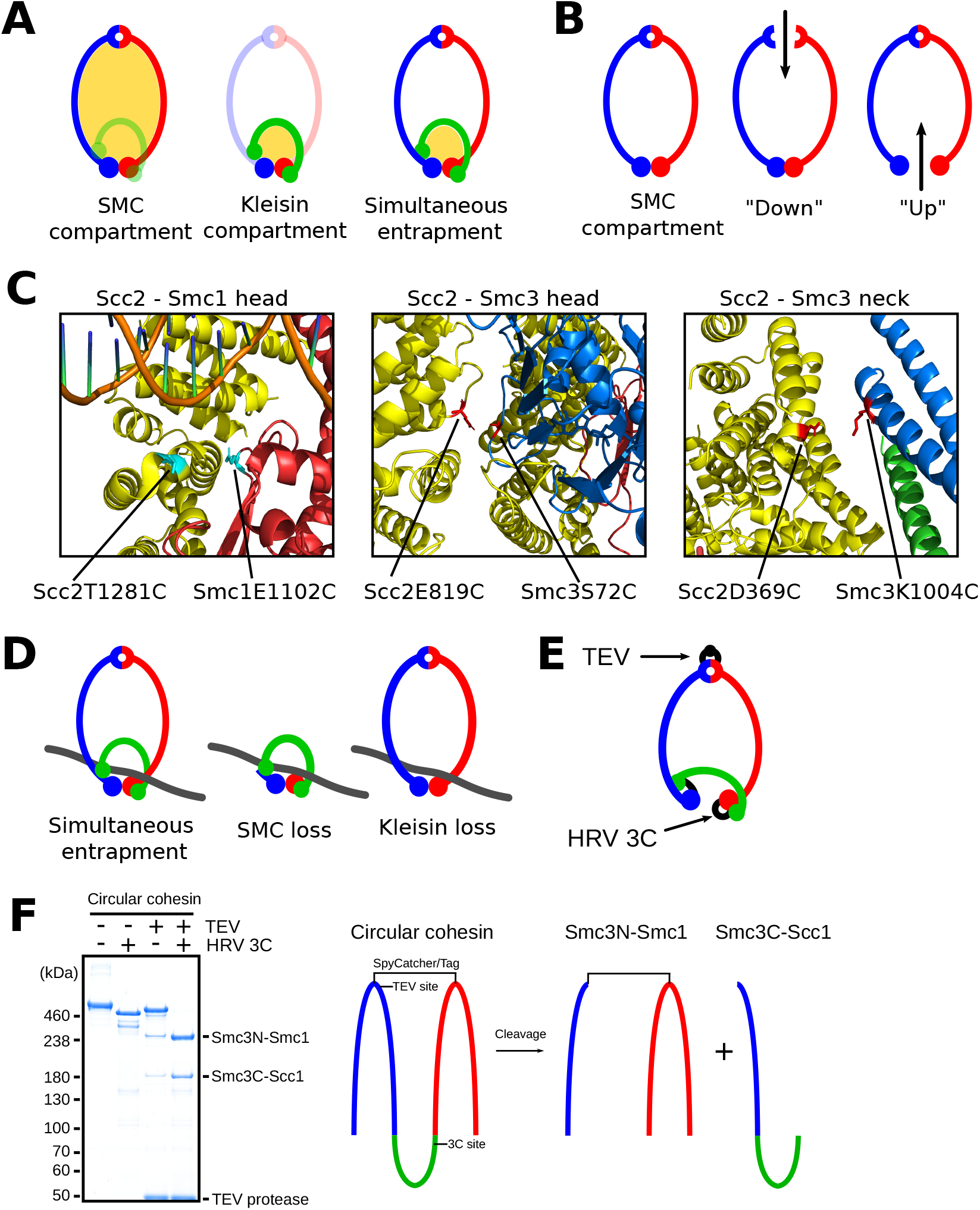
DNA passes through cohesin’s ATPase domains. (A) Models showing the SMC compartment, Kleisin compartment and the overlap between SMC and kleisin compartments, as highlighted in yellow. (B) DNA could enter the SMC compartment by going “down” through the hinge, or “up” through the heads. (C) Left panel showing the cysteine crosslinking pair to crosslink Scc2 to the Smc1 head. Middle panel showing the cysteine crosslinking pair to crosslink Scc2 to the Smc3 head. Right panel showing the cysteine crosslinking pair to crosslink Scc2 to the Smc3 neck. (D) Entrapment through the heads would lead to entrapment in both the SMC and kleisin compartments. If DNA were lost from the SMC compartment, it would still be entrapped in the kleisin compartment, and vice versa. (E) A TEV protease site is located in the linker between the spycatcher and the C-terminal half of Smc3. A HRV 3C protease site is located in the linker between Scc1 and Smc1. (F) Left panel; Coomassie stain of covalently circular cohesin incubated with either TEV and/or HRV 3C proteases for 1 hour at 30 °C in the combinations indicated. Right panel; Schematic showing the polypeptide topology of covalently circular cohesin and the position of the protease cleavage sites. Incubation with both TEV and HRV 3C proteases will cleave the construct into a ~180 kDa C-terminal-Smc3-Scc1 fragment and an N-terminal-Smc3-Smc1 fragment.

We next created two different cysteine pair combinations to measure clamping using our DNA entrapment assay. The first combined Scc2D369C Smc3K1004C with Scc2E819C Smc3S72C, whose simultaneous crosslinking should entrap DNA in a covalent compartment formed by crosslinks between the N-terminal and central domains of Scc2 with Smc3’s neck and head respectively (the clamp; Figure 3D). The second combined Scc2D369C Smc3K1004 and Scc2T1281C Smc1E1102C with Smc1N1192C Smc3R1222C, a pair specific for engaged heads. Simultaneous crosslinking of all three interfaces should entrap DNA in a compartment created by Scc2’s association with both Smc1 and Smc3 heads when they are engaged (Below the clamp or B-clamp; Figure 3D). Under the same conditions that promote entrapment in the SMC compartment, namely the presence of Scc2 and ATP, and the absence of Scc3, DNA was efficiently entrapped in both clamp and B-clamp compartments within 2 min (Figure 3D & E).

### Entrapping DNA in the SMC and kleisin compartments involves passing DNA between ATPase head domains

DNA could enter the SMC compartment by passage “down” through an opened hinge or “up” between SMC heads (Figure 3 – supplement 3B). To distinguish these, we analysed the effect of pre-sealing the hinge interface. This had no effect on SMC entrapment, excluding the possibility that DNA passes “down” through the hinge (Figure 3F). If DNA instead passes between the ATPase heads, without any dissociation of Scc1 from either the Smc3 neck or Smc1 head, then entrapment within the SMC compartment will be accompanied by entrapment between engaged heads and the kleisin subunit associated with them; i.e. in the kleisin compartment, which we have previously shown (Collier et al. 2020). However, it could be argued that entrapment in the kleisin compartment does not arise in this manner but rather as a result of a separate transport process in which DNA passes between a transiently opened SMC/kleisin interface either before or during head engagement. Indeed, such a mechanism has been invoked to explain clamping of DNA on top of engaged heads by Mis4, the *S. pombe* Scc2 ortholog (Higashi et al. 2020). To address whether kleisin disengagement is required for entrapment between engaged heads and their associated kleisin, we introduced the cysteine pair specific for engaged heads (Smc1N1192C Smc3R1222C) into the Smc3-Scc1-Smc1 fusion. Sealing both kleisin interfaces did not prevent entrapment within the kleisin compartment. In fact, this construct entrapped DNA even more efficiently than WT (Figure 3G), presumably because only one single interface needs to be crosslinked compared to the three required for WT. Clearly, entrapment within the kleisin compartment in the presence of Scc2 does not involve passage through either Smc1/ or Smc3/kleisin gates. We conclude that the only interface that must open for entrapment of DNA within the SMC or kleisin compartments is that between the Smc1 and Smc3 ATPase heads.

Due to the identical conditions upon which entrapment in the SMC and kleisin compartments takes place (Collier et al. 2020), it is likely that entrapment in these two compartments is a consequence of a single reaction and that passage of DNA through the head domains leads to simultaneous entrapment of DNA in both the SMC and kleisin compartments (Figure 3 – supplement 3A & D). If true, cohesin with all three interfaces fused should still be able to entrap DNA in the SMC and kleisin compartments. Furthermore, cleavage of either the SMC or kleisin compartments should have no effect on the amount of DNA entrapped, as DNA will remain entrapped within the remaining intact compartment (Figure 3 – supplement 3D). Only simultaneous cleavage of both compartments should release DNA. To test this, we created a version of the covalently circular species of cohesin containing the cysteine pair necessary to crosslink the heads when engaged in the presence of ATP. A pair of tandem TEV protease cleavage sites were inserted in the linker connecting the spycatcher and the Smc3 hinge, which enables cleavage of the SMC compartment, while an HRV 3C protease site was present in the linker connecting Scc1 to Smc1, which enables cleavage of the kleisin compartment (Figure 3 – supplement 3E). Incubating this construct with either TEV or HRV 3C leads to the linearization and opening of the SMC and kleisin compartments, respectively, while incubation with both proteases leads to opening of both compartments, as well as release of a ~180 kDa digestion fragment comprised of the C-terminal half of Smc3 fused to Scc1 (Figure 3 – supplement 3F). Circular cohesin was able to entrap DNA following the BMOE treatment that crosslinks engaged heads (Figure 3H) and remarkably entrapment was largely unaffected by incubation with either TEV or HRV 3C proteases after crosslinking. DNA was only released upon incubation with both proteases. These results imply that DNA is simultaneously entrapped in both the SMC and kleisin compartments due to a single transportation process that involves DNA passage between the Smc1 and Smc3 head domains prior to their engagement. Furthermore, these results allow us to infer the path of Scc1, which must pass over and “above” the DNA, as has been suggested for condensin’s Brn1 subunit (Lee, Rhodes, and Lowe 2022).

## Discussion

In summary, we have shown that cohesin has two DNA gates, one at the hinge and a second at the Smc3/Scc1 interface. Available evidence suggests that the hinge gate is essential for the establishment of sister chromatid cohesion while the Smc3/Scc1 gate is not. Future studies will be required to evaluate whether passage of DNA through the Smc3/Scc1 gate has any in vivo role in addition to releasing cohesin from chromosomes. Passage of DNA through the hinge is likely preceded by and very possibly dependent on its entrapment in a clamp between Scc2 and engaged ATPase heads (Figure 4A), a state created by passage of DNA between the SMC ATPase heads but not through either the hinge or Smc3/Scc1 gate. Following clamping, a section of DNA downstream of the clamp might then be passed through the hinge in a process dependent on Scc3. Passage through the Smc3/Scc1 interface occurs in the absence of Scc2 and may be the result of DNA binding to Scc3 during ATP-driven kleisin disengagement (Figure 4B).

**Figure 4.**
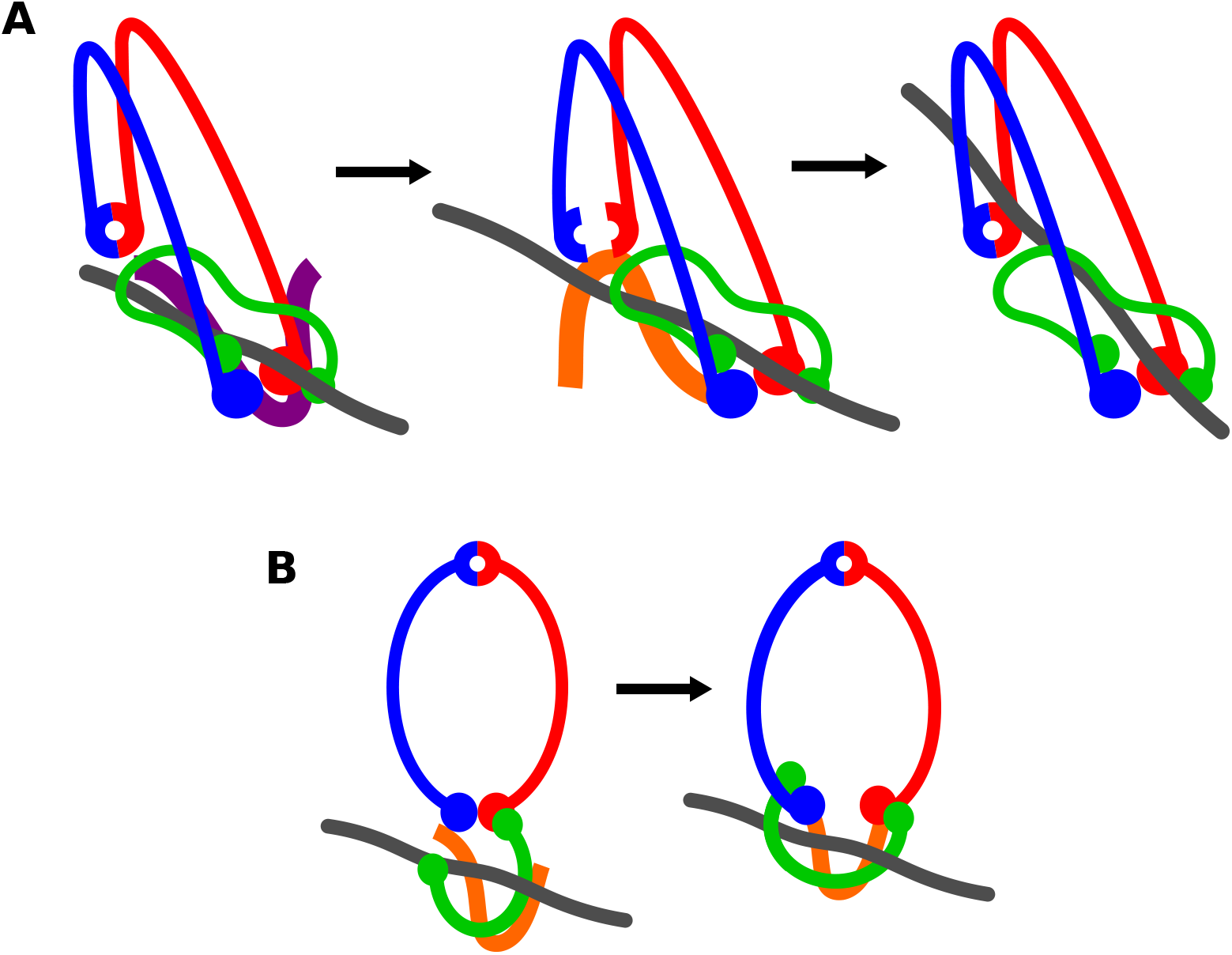
Models for DNA entry. (A) Model for DNA passage through the hinge. Scc2 (purple) first clamps DNA against the Smc3 neck. Scc3 (orange) is involved in opening and passing a downstream section of DNA through the hinge. (B) Model for DNA passage through the Smc3-Scc1 interface. ATP binding leads to the opening of the Smc3-Scc1 interface. DNA then binds to Scc3 and the interfaces closes.

Hitherto, our assay has only detected individual DNAs entrapped inside S-K rings. Entrapment of this nature occurs prior to DNA replication in vivo and is possibly converted to co-entrapment of sister DNAs during S phase with the help of specific replisome proteins (Srinivasan et al. 2020). Crucially, conversion of cohesin that has associated with unreplicated DNA to a form that co-entraps sister DNAs does not require Scc2. If as our in vitro experiments suggest, Scc2 is essential for passage of DNAs through the hinge, then co-entrapment arising during conversion cannot involve any further passage of DNA through the hinge gate. It either involves the Smc3/Scc1 gate (neither Smc3/Scc1 opening nor conversion are essential) or arises from an activity that somehow pulls replicated DNAs through the ring without it being re-opened.

Our assay measuring entry through the hinge in vitro will enable the identification of mutants specifically defective in this process and these can subsequently be used to address whether passage of DNA through the hinge has roles in chromosome topology besides cohesion establishment, for example, in holding together TAD boundaries associated with convergent CTCF sites (Liu and Dekker 2021). SMC hinge domains have two interfaces (north and south) and their dimerization creates a toroidal structure with a narrow lumen that is invariably positively charged (Kurze et al. 2011). Opening, either at one or both north and south interfaces (Gruber et al. 2006; Shi et al. 2020) would enable DNA to bind to highly conserved lysines residing inside the Smc1 hinge (Srinivasan et al. 2018), and this might be an important intermediate stage of the entrapment process. Whether the hinges of SMC complexes besides cohesin also act as DNA gates or whether their positively charged lumens merely bind DNA without passing it inside the ring is an important open question.

## Materials and Methods

### Reagents

|  |  |
| --- | --- |
| ATP (Sigma) | Cat # 11140965001 |
| BMOE (Thermo Scientific) | Cat # 22323 |
| HRV 3C protease (Pierce) | Cat # 88946 |
| TEV protease (Invitrogen) | Cat # 12575023 |
| EnChek phosphate assay kit (Invitrogen) | Cat # E6646 |

### Plasmids

|  |  |
| --- | --- |
| pJC 93 | pACEbac1 Smc1 Smc3 Scc1-StrepII |
| pJC 95 | pACEbac1 Smc1 G22C K639C Smc3 E570C S1043C Scc1-StrepII A547C |
| pJC 153 | pACEbac1 Smc1 K639C N1192C Smc3 E570C R1222C Scc1-StrepII |
| pJC 154 | pACEbac1 Smc1 G22C N1192C Smc3 S1043C R1222C Scc1-StrepII A547C |
| pJC 127 | pACEbac1 Smc1 G22C K639C Smc3Scc1-StrepII E570C A547C |
| pJC 123 | pACEbac1 Smc3 E570C S1043C Scc1Smc1-StrepII K639C |
| pJC 106 | pACEbac1 Smc1 G22C L597ST Smc3 S1043C S606SC Scc1-StrepII A547C |
| pJC 132 | pACEbac1 Smc3Scc1Smc1-StrepII E570C K639C |
| pJC 129 | pACEbac1 Smc1 G22C L597ST Smc3Scc1-StrepII S606SC A547C |
| pJC 124 | pACEbac1 Smc3 S1043C S606SC Scc1Smc1-StrepII ST |
| pJC 135 | pACEbac1 Smc3Scc1Smc1-StrepII S606SC TEV R1222C L597ST N1192C |
| pJC 151 | pACEbac1i Smc1-His E1102C Smc3 Scc1-StrepII |
| pJC 100 | pACEbac1i Smc1 Smc3 S72C Scc1-StrepII |
| pJC 101 | pACEbac1i Smc3 K1004C Smc1 Scc1-StrepII |
| pJC 102 | pACEbac1i Smc1 Smc3 S72C K1004C Scc1-StrepII |
| pJC 105 | pACEbac1i Smc1 E1102C N1192C Smc3 K1004C R1222C Scc1-StrepII |
| pJC 69 | pACEbac1 dN132-Scc2-StrepII |
| pJC 75 | pACEbac1 dN132-Scc2-StrepII T1281C |
| pJC 77 | pACEbac1 dN132-Scc2-StrepII E819C |
| pJC 76 | pACEbac1 dN132 Scc2-StrepII D369C |
| pJC 78 | pACEbac1 dN132-Scc2-StrepII D369C E819C |
| pJC 80 | pACEbac1 dN132-Scc2-StrepII D369C T1281C |
| pJC 84 | pACEbac1 StrepII-Scc3 |

### DNA and protein preparation

Protein and DNA components were prepared as described in Collier et al. 2020.

### ATPase assay

DNA was prepared in DNA buffer (25 mM HEPES pH 7.5, 1 mM TCEP and 5% glycerol) by annealing two complementary single-stranded 40 bp oligonucleotides by heating to 95 °C for 5 min and decreasing in 0.1 °C intervals every 15 s to a final temperature of 4 °C. 150 μl reactions were prepared containing 50 nM cohesin (Smc1, Smc3, Scc1), Scc3, Scc2, and 600 nM DNA in loading buffer (25 mM HEPES pH 7.5, 50 mM NaCl, 1 mM MgCl2, 1 mM TCEP and 5% glycerol), and EnzChek phosphate assay kit components (Invitrogen) added to their recommended concentrations. Reactions were initiated by the addition of ATP (Sigma) to a concentration of 1 mM. The ATPase reaction was followed by measuring the increase in absorbance at 360 nm over 60 min. Data shown an average of three experiments.

### DNA entrapment assay

13 μl reactions were prepared containing 165 nM cohesin and 9.3 nM supercoiled pUC19 in loading buffer (25 mM HEPES pH 7.5, 50 mM NaCl, 1 mM MgCl2, 1 mM TCEP and 5% glycerol). When present, Scc3 was added to a concentration of 165 nM and Scc2 to a concentration of 55 nM. Reactions were initiated by the addition ATP (Sigma) to a concentration of 5 mM. Reactions were incubated at 24 °C for either 40 min or 2 min at 750 rpm. Crosslinking was carried out by addition of 1.5 μl BMOE (Thermo Scientific) to a concentration of 0.64 mM and incubated on ice for 6 min. Samples were then denatured by addition of 1.5 μl 10 % SDS and then incubated at 70 °C for 20 min at 750 rpm. DNA loading dye was then added and samples separated by agarose gel electrophoresis at 50 V for 17 hours at 4 °C. Assays were repeated at least twice.

When cleaving the SMC and kleisin compartments of circular cohesin 10 mM DTT was added after BMOE crosslinking and the samples incubated at 24 °C for 5 min. Protein cleavage was carried out by addition of 1 μl TEV protease (Invitrogen) to cleave the SMC compartment and 1 μl HRV 3C protease (Pierce) to cleave the kleisin compartment. To samples in which one or both proteases was omitted, loading buffer was added instead. Samples were then treated as in other experiments.

### Protein crosslinking assay

10 μl reactions were prepared containing 0.7 μM cohesin (Smc1, Smc3 and Scc1) and Scc2 in loading buffer (25 mM HEPES pH 7.5, 50 mM NaCl, 1 mM MgCl_2_, 1 mM TCEP and 5% glycerol). When present, ATP (Sigma) was added to a concentration of 10 mM and DNA (supercoiled pUC19) added to a concentration of 60 nM. Reactions were incubated at 24 °C for 5 min and then either 1 μl DMSO added, or 1 μl BMOE (Thermo Scientific) added to a concentration of 0.64 mM. Samples were then denatured by addition of 4x LDS protein loading dye and heated at 70 °C for 10 min. Samples were then separated by SDS-PAGE using 3-8 % tris-acetate gels ran at 100 V for 4 hr 30 min at 4 °C.

### Circular cohesin cleavage

10 μl samples were prepared containing 0.7 μM circular cohesin. Protein cleavage was carried out by addition of 1 μL TEV protease (Invitrogen) to cleave the SMC compartment and 1 μL HRV 3C protease (Pierce) to cleave the kleisin compartment. To samples in which one or both proteases was omitted, loading buffer was added instead. Samples were then denatured by addition of 4x LDS protein loading dye and heated at 70 °C for 10 min. Samples were then separated by SDS-PAGE using 3-8 % tris-acetate gels ran at 100 V for 4 hr 30 min at 4 °C.

